# Alpha-1 Antitrypsin Antisense Oligonucleotide Modulates Protease-Antiprotease Imbalance without Further Aggravating Smoke-induced Lung Injury

**DOI:** 10.1101/719682

**Authors:** Rui Xiao, Kyle Stearns, Tina Zelonina, Monica Goldklang, Keith Blomenkamp, Jeffery Teckman, Jeanine D’Armiento

## Abstract

Alpha-1 antitrypsin (AAT) is a serum protease inhibitor that prevents lung injury from protease production during cigarette smoking but causes severe liver disease once mutated. A custom AAT antisense oligonucleotide (ASO) was found to be beneficial for the AATD liver disease by blocking the mutated AAT transcripts. Here we hypothesized that knock-down of AAT aggravates murine lung injury during smoke exposure and acute exacerbations of chronic obstructive pulmonary disease (COPD). C57BL/6J mice were randomly divided into 4 groups for each of the injury models, smoking (inhale for 3 months at 150mg/m^3^) and smoke-flu (inhale for 2 weeks and intranasal influenza virus). The ASO and control (No-ASO) were injected subcutaneously at 10ml/kg of body weight, starting with smoking or four days prior to influenza infection weekly at 50mg/kg. ASO treatment during a 3 month smoke exposure significantly increased the expression of Cela1 mRNA and decreased the serum and lung AAT expression. However, despite the decrease in AAT, neither the inflammatory cell counts in the bronchoalveolar lavage fluid (BALF) nor the lung structural changes were significantly affected by ASO treatment. We observed significant differences in inflammation and emphysema due to smoke exposure alone. With the smoke-flu model, similarly the major differences were found between smoke-flu and room air control, with no additional effect with ASO treatment. Off-target effects or compensatory mechanisms may account for this finding. Alternatively, the reduction of AAT with ASO treatment was not robust enough to lead to lung injury. The result also suggest that the AAT ASO approach for treating liver disease is relatively safe at the specified dose as it did not lead to detrimental outcomes in the lung. These potential mechanisms need to be further investigated in order to fully understand the impact of AAT inhibition on protease-antiprotease imbalance in the murine smoke exposure model.

## Introduction

Alpha-1 antitrypsin (AAT) is a serum protease inhibitor that targets the proteases produced during injury that, in the lung, are ultimately responsible for structural destruction. The protease-antiprotease paradigm has long been recognized and earlier research has identified neutrophil elastase as one of the most potent proteases that mediates destruction in the lung [1, 2]. Alpha-1 antitrypsin inhibits neutrophil elastase activity [3] and prevents the degradation of elastin [4], which helps to maintain the integrity of extra-cellular matrix in the lung. Imbalances between tissue damaging proteases and their inhibitors such as AAT leads to lung destruction and ultimately the development of chronic obstructive pulmonary disease (COPD) [5].

AAT deficiency (AATD) was the first identified genetic predisposition to COPD [6]. Broad screening studies decades ago suggest 2% of patients with COPD have AATD [7]. AATD carriers (Pi*MZ) can account for as high as 17.8% among patient population [8]. AATD has several clinical manifestations; COPD is attributed to a loss of anti-protease activity, and cirrhosis that is attributed to accumulation of misfolded Z-AAT protein within the hepatocyte [9].

Unlike the single gene SERPINA1 that encodes AAT in human, there are five genes (Serpina1a~f) that encode AAT in the mouse. Although a recent study illustrated a mouse model of AATD by CRISPR editing of all five murine AAT genes [10], an antisense oligonucleotide (ASO) approach would still provide insights for the short-term effect and dose dependent responses by targeting the common coding sequences. This ASO approach already demonstrated efficient reduction of human AAT transcript and successfully attenuated the liver injuries associated with AATD [11].

Although beneficial effects were found with AAT ASO treatment on liver disease, the knock down of normal AAT may prove to be detrimental for the lung in the event of an injury with the overproduction of proteases, including chymotrypsin-like elastase 1 (Cela1), a digestive protease that is expressed during lung development [12]. As the main etiology of COPD, smoking induces inflammation and tips the balance of protease/antiprotease balance to increased proteases activity and therefore lead to a distinct gene expression profile [13] and subsequently parenchymal destruction [14–16]. Another main trigger for rapid progression of COPD is respiratory viral or bacterial infection [17] and in particular viruses, such as picornavirus and influenza, are among the highest prevalence in patients with acute exacerbation of COPD [18].

Here we hypothesize that murine AAT ASO treatment decreases the expression of AAT and subsequently aggravate the smoke-induced lung injury and acute exacerbation modeled by influenza virus infection after smoke exposure.

With the custom designed mouse AAT ASO, we can understand if AAT ASO inhibits the expression of AAT in the lung similar to that in the liver. In addition, we can determine if AAT modulation affects the outcome of smoking-induced lung injury and acute COPD exacerbation in assessing the overall feasibility for using AAT ASO for treatment of liver disease in patients with a pre-disposition to proteolytic lung destruction.

## Materials and Methods

### Alpha-1 Antitrypsin Antisense Oligonucleotide Treatment

Mouse AAT ASO and control ASO solution were prepared by Ionis Pharmaceutical in collaboration with Keith Blomenkamp and Dr. Jeff Teckman in Saint Louis University with the method described previously [11]. Two filter flasks were aliquoted upon arrival at 5mg/ml concentration and administered via subcutaneous injection with a dose of 10ml/kg body weight once a week, to make up a final weekly concentration of 50mg/kg.

### Animal housing and smoke exposure

60 male C57BL/6J mice, 10-12 weeks of age, purchased from Jackson Laboratory (Bar Harbor, ME), were randomly divided into 4 groups, Air-NoASO (AN), Air-ASO (AA), Smoke-NoASO (SN), and Smoke-ASO (SA). All mice were acclimated to the animal facility for at least 48 hours prior to use and were given chow diet and water ad libitum. Smoke exposure was conducted on TE-10 Teague Smoking Apparatus (Teague Enterprises, Woodland, CA) with 3R4F Reference Cigarettes (University of Kentucky, Lexington, KY) for 5 days per week, 5 hours per day for a total of 3 months at a total particulate matter of 100~150mg/m^3^. Total particulate matter was monitored by aerosol monitor DustTrak II 8530 (TSI, Shoreview, MN) and confirmed by gravimetric analysis. After starting the smoke exposure, alpha-1 antitrypsin antisense oligonucleotide treatment was administered weekly (Figure 1).

**Figure 1.**
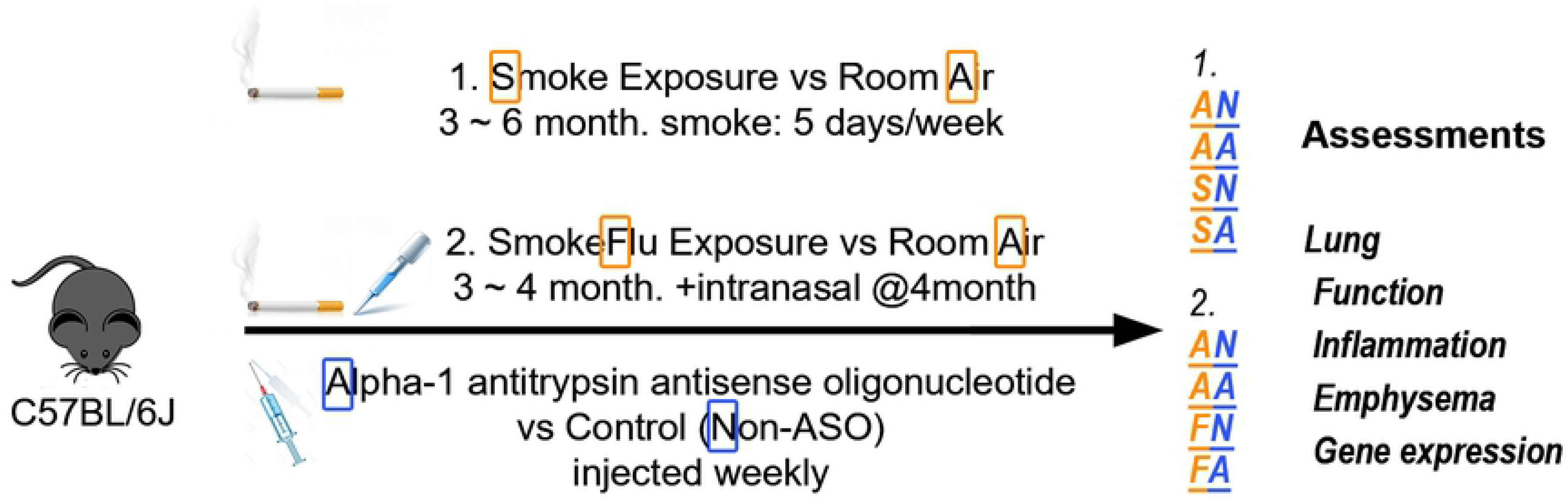
Experimental design. Alpha-1 antitrypsin antisense oligonucleotide treatment was tested on two sets of lung injury models, smoke exposure for chronic obstructive pulmonary disease (COPD) and additional flu infection for COPD exacerbation. Each experiment was randomly divided into 4 groups, first letter indicating the lung injury model, second letter whether treated with ASO.

### Influenza infection model

22 male C57BL/6J mice, 10-12 weeks of age, from Jackson Laboratory (Bar Harbor, ME), were randomly divided into 4 groups for another set of comparisons, Air-NoASO (AN), Air-ASO (AA), SmokeFlu-NoASO (FN), and SmokeFlu-ASO (FA). After smoke exposure for two weeks as described above, mice were anesthetized with isoflurane (1-5%) and infected intranasally with a dose of 1000 TCID50 Influenza A/Puerto Rico/8/1934 H1N1 virus suspension in 30µl of PBS. All laboratory personnel who perform infections and handle infected mice have undergone training for BSL-2 procedures to enter and use animals in this facility. Two oligonucleotide doses were given 1 and 4 days prior to influenza infection and continued once a week post-infection (Figure 1). As the infection dose may still result in animal death, starting from the first antisense treatment which is 4 days before the infection, animals were weighted and monitored for health on a daily basis by qualitative assessment of general health noting clinical signs of morbidity such as ruffled fur, hunched posture and shivering. The humane endpoint was defined as 20% weight loss and appear moribund as assessed by lack of movement, hunched posture, and unkempt appearance. If within 24 hours mice did not recover and regain weight, they were euthanized. At two weeks after the infection, 4 out of 8 SmokeFlu-NoASO (FN) mice and 3 out of 6 SmokeFlu-ASO (FA) mice died abruptly before reaching pre-defined humane endpoint. The control groups, 3 Air-NoASO (AN) and 5 Air-ASO (AA) did not suffer from significant weight loss or adverse effects. Animals are handled exclusively in biosafety cabinets within the animal facility, including anesthetizing, infecting, and sacrificing of mice, and surfaces will be cleaned with antiseptic solution provided.

### Lung Function Testing

We assessed the lung function of 60 mice for smoke-antisense study. After 3 months of smoke exposure and weekly injections, lung function was measured with flexiVent FX2 (EMKA, Montréal, QC, Canada), a computer-controlled piston ventilator that measures lung function in small animals. The mice were sedated with pentobarbital 75 mg/kg, tracheostomized and intubated with an 18-gauge beveled tracheal tube to connect directly to flexivVent. After 3–5 min equilibration on the ventilator with a tidal volume of 8 ml/kg and frequency of 150 breaths/min and maintaining the mice at 37°C with a homeothermic blanket (Homeothermic Blanket System; Harvard Apparatus, Holliston, MA), mice were given succinylcholine 0.5 mg every 14 min by i.p. injection. Pertubations, including deep inflation, snapshot-150, quick prime-3, and PV loop (PVs-P) were performed using the flexiVent system.

### Inflammation and emphysema assessment of the lung

For both smoke-antisense and smoke-flu-antisense studies, lungs were lavaged during sacrifice once with 500µl of PBS to obtain concentrated BAL supernatant, followed by two more lavages with 1 ml of sterile PBS to maximize cell yields. Total cells were counted with a TC20 automatic cell counter (Bio-Rad, Philadelphia, PA). Cytospin preparations were stained with Quick-Diff (Imeb), and cells were analyzed for differential counts using morphological criteria. Left lung was formalin-fixed, paraffin-embedded, step-sectioned every 200µm and stained by H&E method. Morphometric analyses, including the traditional gridline counting and parenchymal airspace profiling methods [19], were performed on the lung sections.

### Western Blotting, ELISA, and qRT-PCR

The AAT protein expression in the lung was assessed with western blotting (alpha-1 antitrypsin antibody, #16382-1-AP, rabbit polyclonal, Proteintech, Rosemont, IL). Serum level of AAT was evaluated with Mouse Alpha 1 Antitrypsin ELISA Kit (Product: ab205088, Let Number: GR3272327). PCR probe for Cela1 (chymotrypsin-like elastase family, member 1) was purchased from ThermoFisher (Applied Biosystems, Foster City, CA; Assay ID: Mm00712898_m1; Cat #4331182). Cela1 was implicated with mouse lung development and human AATD [12, 20].

### Statistical analysis

Simple pairwise comparisons were performed with Students t-test or Wilcoxon-test (non-parametric test for the survival curve) assuming unequal variance. Additional tests were performed after grouping both ASO control and treated mice, for example, overall flu (FN&FA) vs control (AN&AA), when ASO treatment did not make any significant difference. All error bars indicate mean ± SEM.

## Results

### ASO treatment effectively knocked down the AAT protein expression and induced the expression of Cela1 in the lung

Endogenous AAT protein expression in the serum (Figure 2A, P<0.005; N=8~10) and the lung (Figure 2B, P<0.001; N=6, from 3 western blots, Figure S2) and were significantly reduced in mice injected with ASO. The injection also significantly increased the Cela1 expression for the air-exposed mice (AA vs AN, P=0.004) but not for the smoke-exposed mice (SA vs SN, P=0.8; Figure 2C).

**Figure 2.**
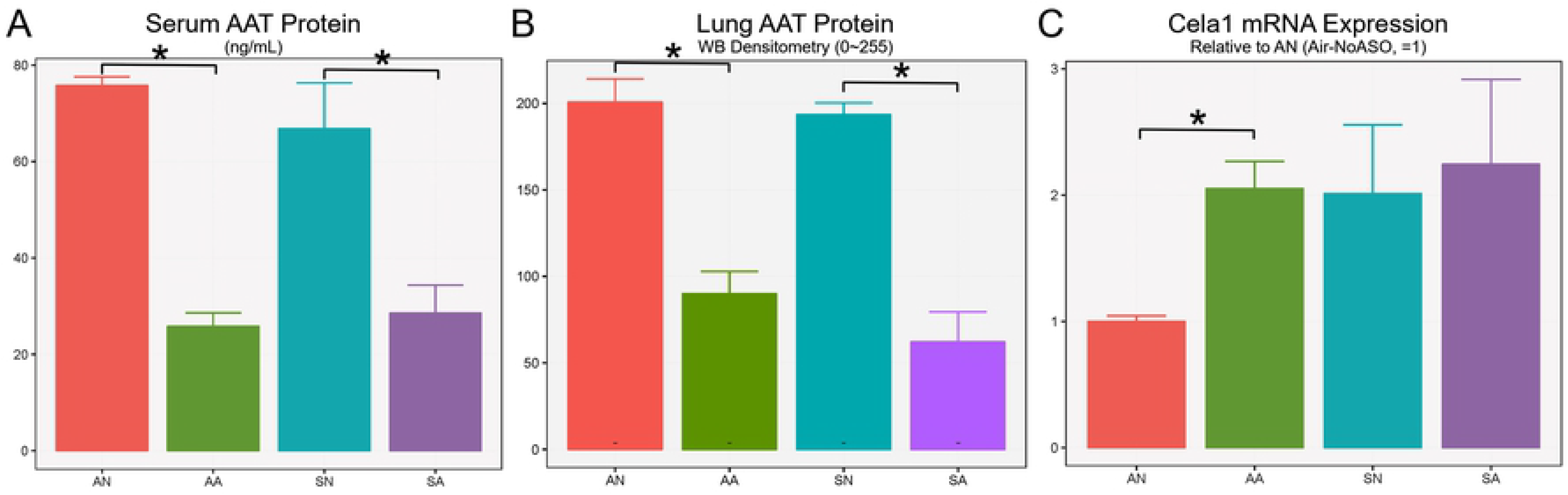
Alpha-1 antitrypsin (AAT) expression was significantly decreased by the antisense oligonucleotide treatment. Although the treatment lead to decreased AAT in the serum (A; P<0.005) and the lung (B; P<0.001) and increased Cela1 elastase expression (C; P=0.014), Cela1 was highly expressed in smoke exposed mice and did not increase further by oligonucleotide treatment (P=0.80).

### ASO did not significantly modulate lung responses of mice in a smoke-induced lung injury model

Lung function testing results measured by flexiVent showed an overall significantly increased inspiratory capacity for mice exposed to smoke (SN&SA vs AN&AA, P=0.01, Figure 3A). ASO treatment did not contribute to any changes in inspiratory capacity, resistance, or compliance regardless of the exposure conditions (Figure 3B).

**Figure 3.**
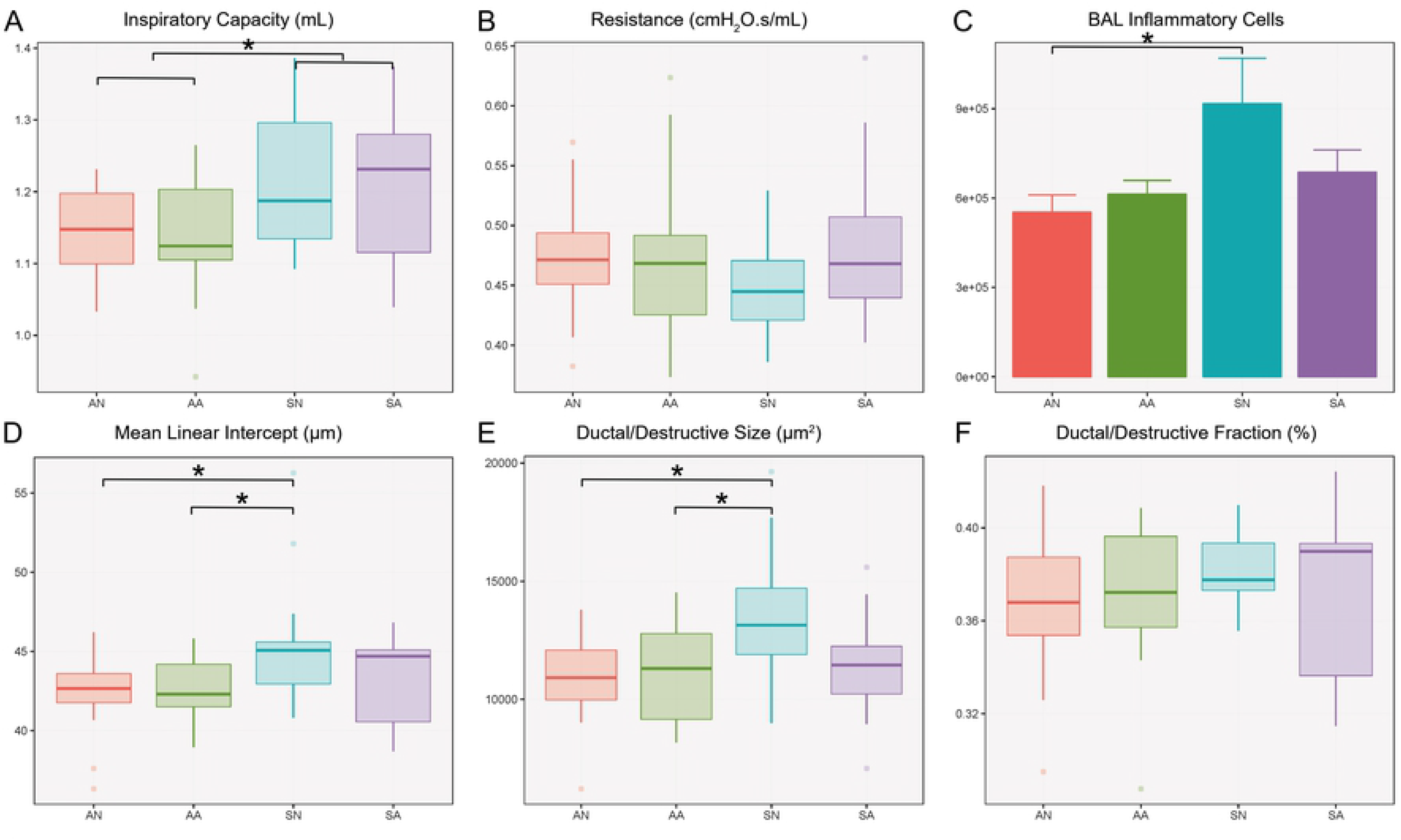
In the COPD model built on 3-month of smoke exposure, smoke exposure contributes to changes in inspiratory capacity (A; SN&SA vs AN&AA, P=0.01), inflammation (C; bronchoalveolar lavage cell counts, P=0.04) and emphysema (D-F; mean linear intercept, P=0.023; ductal/destructive size, P=0.008) but antisense treatment did not make any changes in lung function, inflammation, or emphysema

Despite the significantly increased inflammatory cells in the bronchoalveolar lavage fluid (BALF) of the smoke-exposed mice (SN vs AN, P=0.04), the BALF inflammatory cell counts were not significantly affected by ASO treatment (P>0.19; Figure 3C). Analysis of the lung morphometry in the mice demonstrated emphysema development with smoke exposure alone (Mean Linear Intercept, P=0.02; Ductal/destructive Size, P=0.008), which was not further aggravated by the reduction of AAT with the ASO (P>0.05). Although the ASO treatment successfully decreased the expression of AAT in the serum and the lung, no additive effect was seen when these mice were exposed to smoke (Figure 3A~3F).

### ASO did not significantly affect survival and lung responses of mice exposed to smoke and influenza infection

Both groups exposed to smoke and influenza virus exhibited mortality of ~50% two weeks after the infection (Figure 4A), which was also accompanied by significant weight loss. The most weight loss (26.49%) was observed at day 12 post infection followed by weight gain until sacrifice date (Figure 4B).

**Figure 4.**
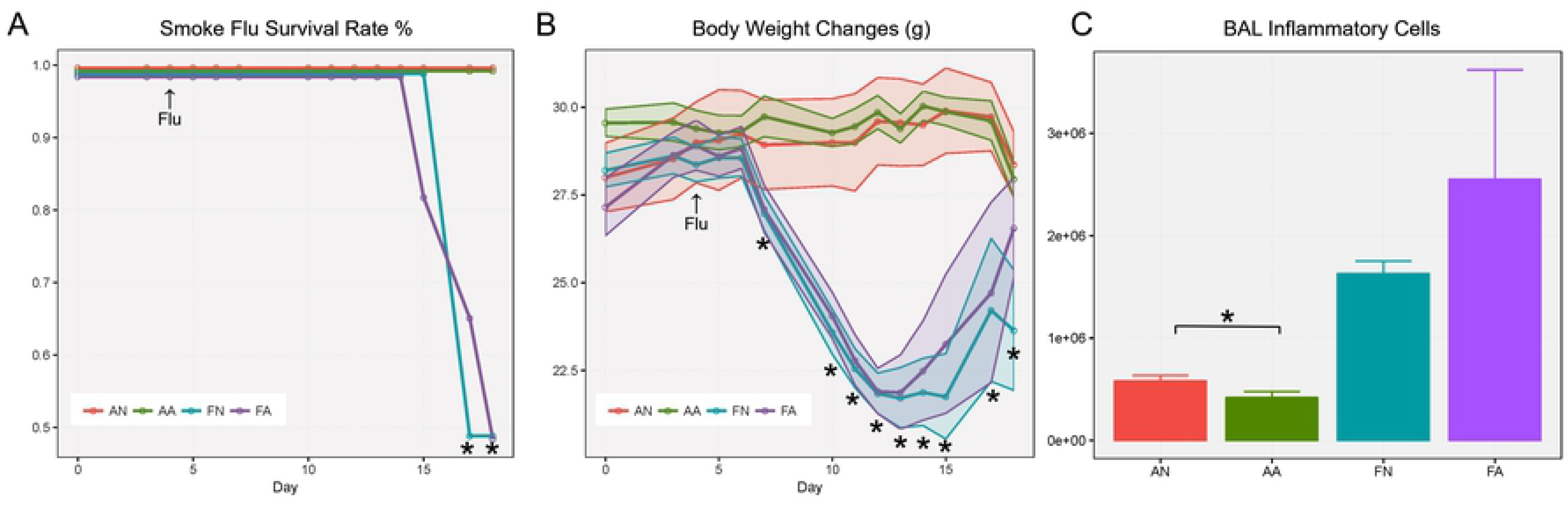
With the COPD exacerbation model, the survival rate (A) and weight change (B) were only affected by smoke-flu injury but not antisense treatment (up to day 18, FA&FA vs AA&AN, P<0.05). Inflammatory cells were also primarily affected by smoke-flu injury, antisense treatment made minor differences to mice without injury (C; AA vs AN, P=0.015) but still did not modulate the responses in the smoke-flu injured mice (FA vs FN, P=0.93).

Similar to smoke exposure alone, there was significantly increased inflammation due to smoke exposure and flu infection (FN vs AN, P=0.018). However, the ASO treatment did not contribute to any significant changes (P>0.1). Between all smoke-flu exposed and air exposed mice, there are significant differences at day #17 and #18 after first dose of antisense oligonucleotide for the survival rate and between day #7 and day #18 for the body weight.

## Discussion

Consistent with the previous study, where an 80% reduction in circulating AAT was found in the liver after AAT ASO treatment, we also found more than 50% reduction in AAT protein expression in the serum and the lung (Figure 2A-B). As one of the digestive enzymes that can be covalently neutralized by AAT and played an important role in emphysema development, we first examined the expression of chymotrypsin-like elastase 1 (Cela1) [20]. Although ASO did not further increase the expression of Cela1 for the mice exposed to cigarette smoke, it did bring up Cela1 expression of air-exposed mice to a comparable level to those exposed to smoke. This suggests that the smoke exposure by itself may have already maximized Cela1 expression. Therefore, down-regulating protease inhibitor would not favor further over-expression of Cela (Figure 2C). From another perspective, the AAT expression is still present at a detectable level (Figure 2A-B). This level of AAT may remain effective and functional, especially in comparison to a recently published paper showing complete knock-out of AAT resulted in significant changes in lung function and emphysema [10].

In the initial study built upon an animal model of COPD, even with a sample size of more than 10 per group, we did not observe any significant differences attributed by ASO treatment. This includes lung functional parameters, inflammatory cells, and lung structural changes. As expected the differences between smoke alone and air control can be found in degrees of inflammation and emphysema. However, other than noticeable weight loss in the ASO treated mice (Figure S1), which is also reported to be a side effect when used in neurodegenerative disorders [21], partial knockdown of AAT is unable to modulate inflammatory and emphysematous changes in the lung (Figure 3).

To examine whether partial knockdown of AAT plays any role in COPD exacerbation, we designed another set of experiments that utilize both smoke exposure and influenza virus. The exacerbation model itself clearly distinguished the mice exposed to smoke and influenza from room-air control even with limited survival rate (~50%). However, the ASO treatment did not make any significant differences (FA vs FN) for mice exposed to both smoke and flu, which are the most important groups that may lead to therapeutic discoveries.

In summary, ASO do present an effective method to knock down AAT expression and a valuable vehicle for mechanistic studies regarding AAT mediated protease/antiprotease balance. However, in both scenarios that we investigated, AAT modulation did not significantly alter the outcome of smoke-induced lung injury and additional viral infection. Considering the Cela1 expression, we suspect that the main explanation is that the protease expression was fully saturated in either model and therefore unresponsive to the decreased AAT. Another explanation is the potential off-target effects and compensatory mechanisms that was triggered due to ASO treatment. The antisense design may target unintended proteins with similar nucleotide sequence. Further, significantly reduced AAT expression was found to up-regulate neutrophil elastase and may impact the expression of other key proteases. These potential mechanisms need to be further investigated in order to fully understand the protease-antiprotease imbalance pertaining to AAT in the murine smoke exposure model.

Most importantly, this study did not demonstrate significant worsening of lung injury in murine models of COPD exacerbations as a result of reduced liver AAT expression. These results suggest that knockdown of AAT expression in PiZZ patients for liver protection may not have detrimental effects on the lung. Therefore the use of AAT ASO treatment in AAT Z-polymer associated liver disease appears to be a viable option, likely in combination with AAT augmentation therapy for individuals at risk for COPD development.

## Acknowledgements

We thank former D’Armiento laboratory member Dr. Emilio Arteaga-Solis for insightful discussions.

All animal studies were performed with the approval of the Institutional Animal Care and Use Committee of Columbia University.

This work was supported by National Institutes of Health/National Heart, Lung, and Blood Institute grant R01HL086936 (J.M.D.), American Lung Association/Alpha-1 Foundation research grant RG-361233 (R.X.) and The Center for LAM and Rare Lung Disease at Columbia University.

The authors declare that they have no conflicts of interest.

RX, MG, and JMD designed the study. KB and JT provided the antisense for the experiment. RX, KS, TZ and MG performed the experiments. All authors read and approved the final manuscript.

## Supporting Information

**Figure S1.**
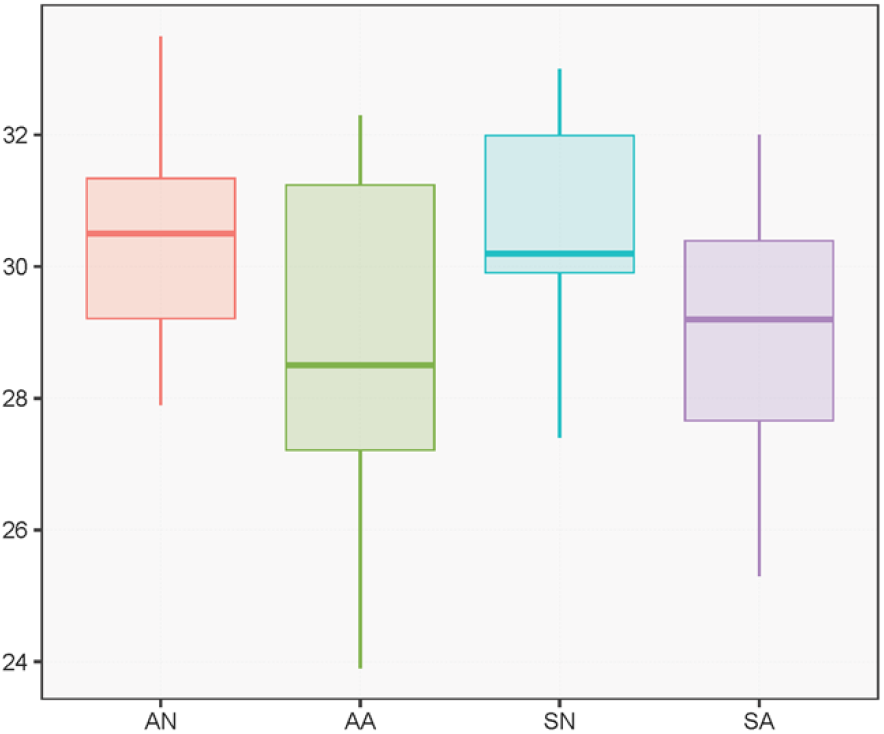
Alpha-1 antitrypsin antisense oligonucleotide treatment resulted in decreased body weight in mice (AA vs AN, SA vs SN, P<0.1; AA&SA vs AN&SN, P=0.01).

**Figure S2.**
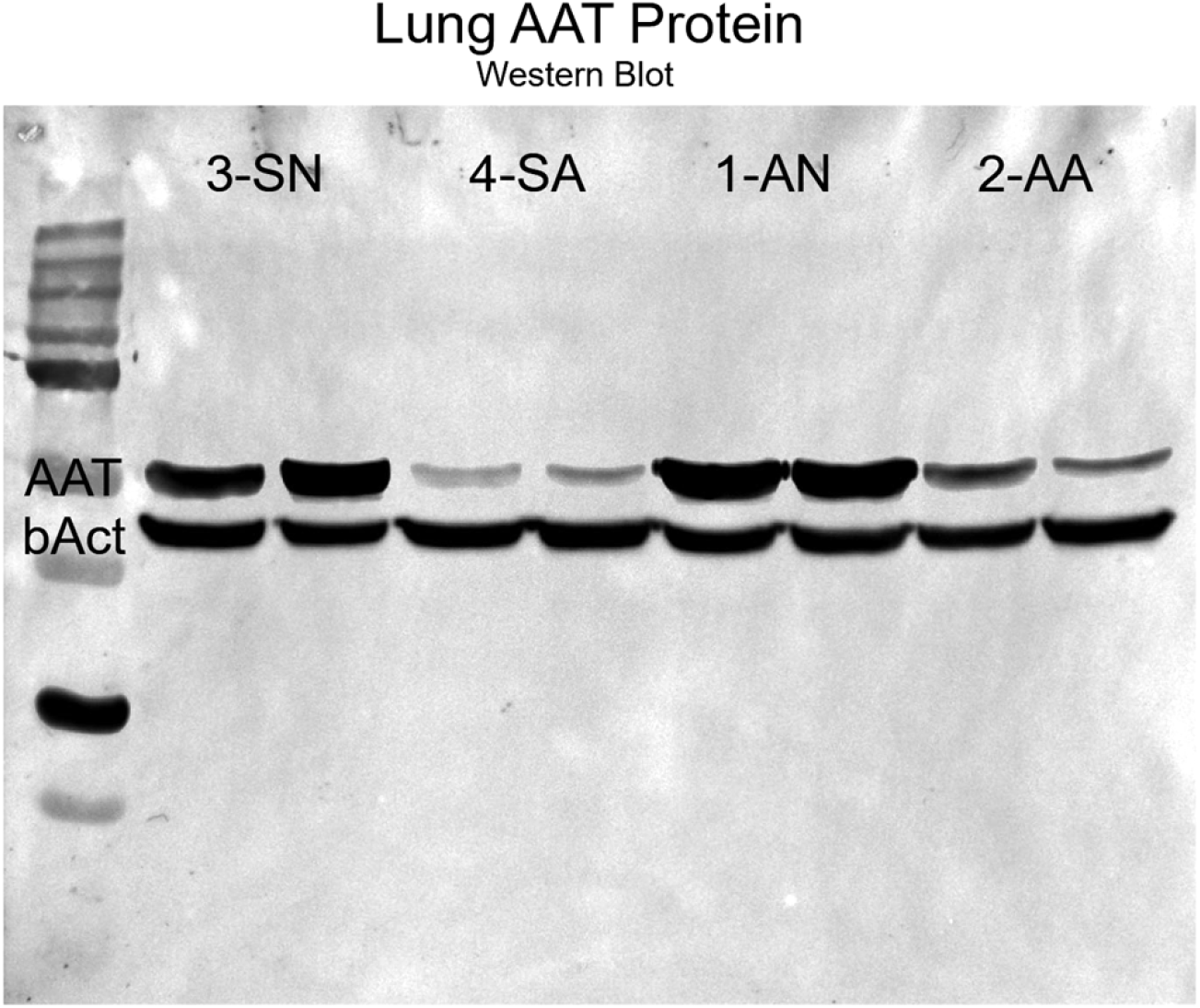
Alpha-1 antitrypsin antisense oligonucleotide treatment resulted in decreased AAT expression in the lung, one representative western blot (N=2) contributing to Figure 2B.

